# Direct determination of diploid genome sequences

**DOI:** 10.1101/070425

**Authors:** Neil I. Weisenfeld, Vijay Kumar, Preyas Shah, Deanna M. Church, David B. Jaffe

## Abstract

Determining the genome sequence of an organism is challenging, yet fundamental to understanding its biology. Over the past decade, thousands of human genomes have been sequenced, contributing deeply to biomedical research. In the vast majority of cases, these have been analyzed by aligning sequence reads to a single reference genome, biasing the resulting analyses and, in general, failing to capture sequences novel to a given genome.

Some *de novo* assemblies have been constructed, free of reference bias, but nearly all were constructed by merging homologous loci into single ‘consensus’ sequences, generally absent from nature. These assemblies do not correctly represent the diploid biology of an individual. In exactly two cases, true diploid *de novo* assemblies have been made, at great expense. One was generated using Sanger sequencing and one using thousands of clone pools.

Here we demonstrate a straightforward and low-cost method for creating true diploid *de novo* assemblies. We make a single library from ~1 ng of high molecular weight DNA, using the 10x Genomics microfluidic platform to partition the genome. We applied this technique to seven human samples, generating low-cost HiSeq X data, then assembled these using a new ‘pushbutton’ algorithm, Supernova. Each computation took two days on a single server. Each yielded contigs longer than 100 kb, phase blocks longer than 2.5 Mb, and scaffolds longer than 15 Mb. Our method provides a scalable capability for determining the actual diploid genome sequence in a sample, opening the door to new approaches in genomic biology and medicine.

---

Determining the genome sequence of an individual organism is of fundamental importance to biology and medicine. While the ability to correlate sequence with specific phenotypes has improved our understanding of human disease, the molecular basis of ~20% of mendelian phenotypes is still unknown (http://omim.org/statistics/geneMap), and the situation for common disease is much worse. Contributing to this is the incomplete elucidation of the genomic architecture of the genomes under study (Eichler et al. 2010).

Decades of research have yielded a vast array of laboratory and computational approaches directed at the problem of knowing the genome sequence of a given sample. These vary dramatically in their aggregate experimental burden, including input DNA amount, organizational complexity, laboratory and computational requirements for expertise and hardware, project complexity, cost and timeline, with greater burden tending to yield a higher quality genome sequence.

At the low end, and by far the most widely executed, are ‘resequencing’ methods that generate short reads, then align them to a haploid reference sequence from the same species, to identify differences with it, thereby partially inferring the sequence of the sample (Li et al. 2008; McKenna et al. 2010). Several projects have generated and analyzed over a thousand human samples each, yielding extraordinarily deep information across populations (1000 Genomes Project Consortium et al. 2015; Gudbjartsson et al. 2015; Nagasaki et al. 2015), although in general such methods cannot completely catalog large-scale changes, nor distinguish between parental alleles. Moreover, such methods are intrinsically biased by comparison to a reference sequence, thus limiting their ability to see sequences in a sample that are significantly different from it. (Chaisson, Huddleston et al. 2015).

By contrast, an analysis of an individual genome would ideally start by reconstructing the genome sequence of the sample, without using a reference sequence. This *de novo* assembly process is difficult for large and complex genomes (Istrail et al. 2004; Chaisson, Wilson et al. 2015; Gordon et al. 2016; Steinberg et al. submitted). A core challenge is the correct representation of highly similar sequences, which range in scale from single base repeats (homopolymers) to large complex events including segmental duplications (Bailey et al. 2002).

There is an even larger scale at which similar sequences appear: homologous chromosomes, which are ‘repeats’ across their entire extent. To correctly understand the biology of a diploid organism, these homologous chromosomes need to be separately represented (or phased), at least at the scale of genes (Tewhey et al. 2011; Muers 2011; Glusman et al. 2014; Snyder et al. 2015). This is required to correctly understand allele-specific expression and compound heterozygosity. For example, two frameshifts in one gene allele could have a completely different phenotype than one each in both alleles; likewise larger-scale effects such as changes to gene copy number (Horton et al. 2008; Pyo et al. 2010) need to be understood separately for each homologous chromosome. However, precisely because homologous chromosomes are so similar, it is challenging to keep them separate in assemblies.

In fact, the standard of the field for genome assembly has been to represent homologous loci by a single haploid ‘consensus’ sequence, that merges parental chromosomes. This loses ‘half’ of the information, and in general does not represent a true physical sequence present in nature. As a step in the right direction, one could generate a haploid assembly together with a phased catalog of differences between the two originating chromosomes (Pendleton et al. 2015; Mostovoy et al. 2016). However for the same reasons that a *de novo* assembly can carry more information than a simple catalog of differences with a reference sequence, these ‘phased haploid’ assemblies carry less information than true diploid assemblies that separately display homologous loci. In a few cases these diploid *de novo* assemblies have been demonstrated for small and mid-sized genomes (Jones et al 2004; Chin et al. 2016). There are two extant instances of diploid *de novo* assemblies of human genomes, one obtained by Sanger sequencing of multiple libraries (Levy et al. 2007), and one from thousands of separate clone pools, each representing a small, low-throughput partition of the genome (Cao et al. 2005).

In this work we bridge the gap between low-cost resequencing approaches, and high-cost diploid assembly approaches, by creating diploid *de novo* assemblies, at very low experimental burden. Our method is also based on genome partitioning. Using an automated microfluidic system (Zheng et al. 2016), we are able to generate the entirety of data for an assembly project from one library. Moreover, this library is made from about one nanogram of high molecular weight DNA, far less than alternative approaches. The cost of our data is in the range of low end methods based on read alignment, and special expertise is not required for assembly, because the process is automatic.

To demonstrate our method, we chose seven human samples from diverse populations, including four for which parental data was available, allowing us to test phasing accuracy. This set also included a sample for which 340 Mb of finished sequence was generated during the Human Genome Project, thus providing an unprecedented source of truth data, and allowing us to assess the fine-scale accuracy of our method, in a way that has not been previously possible. We assembled these samples using a new algorithm called Supernova Assembler. Both the laboratory and computational methods are encapsulated in a complete commercial system from 10x Genomics. Open source software, datasets and assemblies described in this work are publicly available.

## RESULTS

### Data generation

We provide a conceptual explanation of our schema for data generation, including the fundamental characteristics of the data type. Our method uses an updated version of the 10x microfluidic gel bead partitioning system (Zheng et al. 2016), called the Chromium Genome Reagent Kit (**Methods**). Library construction starts from 1.25 ng of DNA having size 50 kb or longer (Zhang et al. 2012), from a single individual organism or clonal population (such as a cell line).

Briefly, the system exploits a reagent consisting of several million gel beads, with each bead containing many copies of a 16-base barcode unique to that bead. A microfluidic device delivers individual beads into approximately one million partitions, along with high-molecular weight genomic DNA molecules and reagents. Each partition receives several long molecules (as discussed below), and the molecular biology of the system is arranged to create constructs having the barcode, along with ~350 bp of genomic DNA from a molecule, sandwiched between Illumina adapters. The barcode is placed at the beginning of the first read in a pair. These constructs are then sequenced on an Illumina instrument, yielding groups of read pairs, organized by barcode. Sets of these read pairs which originate from the same molecule are called Linked-Reads.

We describe the sequencing configuration that we use for *de novo* assembly. Paired reads of length 150 bases each are generated. This read length was chosen so that data could be sequenced on the HiSeq X instrument, which yields the lowest cost data among Illumina instruments, and which has a maximum read length of 150. Data can also be generated on the HiSeq 2500 in rapid run mode. We tested the HiSeq 4000, observing a two-fold reduction in contig size (**Supp. Note 1**). We recommend that samples be sequenced to 56x, or about 1200M reads for a human genome. Lower coverage is possible and described later.

Here we describe the model behavior of the system. Of the DNA that is loaded, approximately 40% makes it completely through the process and contributes to the sequencing library. For 1.25 ng of loaded material, distributed across 10^6^ partitions, and supposing that all molecules had size 50 kb, the mean number of molecules per partition would be about 10, in total representing ~0.5 Mb of genome per partition. At 56x coverage, the mean number of Linked-Reads (read pairs) per molecule for a human genome would thus be (1200M/2) / (10^6^ × 10) = 60, and covering the molecule to depth (120*150) / (50,000) = 0.36x. Importantly, the mode of operation of the system is to provide <u>shallow coverage by many reads for each of many long molecules</u>. In particular the system is not designed to deeply cover individual molecules, in contrast to Synthetic Long-Reads (Voskoboynik et al. 2013).

For smaller genomes, assuming that the same DNA mass was loaded and that the library was sequenced to the same read-depth, the number of Linked-Reads (read pairs) per molecule would drop proportionally, which would reduce the power of the data type. For example, for a genome whose size is 1/10th the size of the human genome (320 Mb), the mean number of Linked-Reads per molecule would be about 6, and the distance between Linked-Reads would be about 8 kb, making it hard to anchor barcodes to short initial contigs. Modifications to workflow, such as loading less DNA and/or increasing coverage would be potential solutions for smaller genomes, but are not described here.

Using the method described above, we generated datasets from seven human individuals of varied ancestry and sex (**Table 1**). All were created from DNA of size > 90 kb, measured by length-weighted mean (**Table 1**). We tested the performance of the system on DNA of several different sizes, noting degradation in performance (**Supp. Note 2**), particularly for DNA < 30 kb. (DNA of size ~20 kb yielded scaffolds of N50 size 0.6 Mb, whereas DNA of size ~50 kb yielded scaffolds of N50 size 12.8 Mb.) We also tested the effect of reducing coverage from 56x to 38x (**Supp. Note 3**), noting some degradation in quality, e.g. scaffold N50 declining from 17 to 12 Mb.

**Table 1.** Genome assemblies. Assemblies of this work plus preexisting assemblies (H from Pendleton et al. 2015, I from Mostovoy et al. 2016, J from Putnam et al. 2016, M from Cao et al. 2015; see **Supp. Note 4**). All statistics were computed after removing scaffolds shorter than 10 kb. Comparisons to reference use GRCh37 (chr1-22, X,Y), with chrY excluded for female samples. **Id:** identifier of assembly in this table. **Sample:** source of starting material. HGP is from the donor to the Human Genome Project for libraries RPCI 1,3,4,5 (http://bacpac.chori.org/library.php?id=1), for which 340 Mb of finished sequence are in GenBank. HGP was from fresh blood, others are Coriell cell lines. **Ethnicity:** ethnicity of individual. **Sex:** sex of individual. **Data description:** capsule description of data type. **X:** estimated coverage of genome by sequence reads. For assemblies of this work, reads were 2x150; 1200M reads were used for each assembly; all samples were sequenced on HiSeq X. **F:** inferred length-weighted mean molecule length of DNA in kb (see **Supp. Table 1** for other statistics). **N50 contig size:** N50 size of FASTA records, after breaking at sequences of 10 or more n or N characters. **N50 phase block size:** N50 size of phase blocks, computed for A-G, and as reported for assembly M. **N50 scaffold size:** N50 size of FASTA records, excluding Ns. **Gappiness:** fraction of bases that are ambiguous. **N50 perfect stretch:** N50 length in kb of segments on finished sequence from same sample that are perfectly mirrored in assembly (see text). **Phasing error rate:** fraction of phased sites in megabubble branches whose phasing did not agree with the majority. **Missing kmers:** fraction of 100-mers in reference that are missing from the assembly (includes *bona fide* sample/reference differences). **Haploid:** value for haploid version of assembly. **Diploid:** value for diploid version of assembly. **Inconsistent at given distance:** of kmer pairs at the given distance in the assembly, and for which both are uniquely placed on the reference, fraction for which either the reference chromosome, orientation, order, or separation (± 10%) are inconsistent (includes *bona fide* sample/reference differences). **Wall clock:** run time (days) for assemblies using a single server having 384 GB available memory (booted with "mem=384G"), exclusive of subsampling to 1200M reads, sorting by barcode and trimming of barcodes (total 2-5 hours). **Software used to create assemblies: [A-G]** Supernova 1.1 with default parameters; **[H]** Falcon, Blasr (Chaisson and Tesler 2012), Celera Assembler (Koren et al. 2012), RefAligner (Nguyen 2010; Anantharaman and Mishra 2001), custom scripts; **[I]** SOAPdenovo2 (Luo et al. 2012), ABYSS (Simpson et al. 2009), Longranger (Zheng et al. 2016), BWA-MEM (Li 2013), fragScaff (Adey et al. 2014), RefAligner, Lastz (Harris 2007), BioNano hybrid scaffold tool (Mak et al. 2016); **[J]** Meraculous (Chapman et al. 2011), HiRise (Putnam et al. 2016); **[K-L]** Celera Assembler, Quiver (Chin et al. 2013); **[M]** SOAPdenovo2, ReFHap (Duitama et al. 2012), custom pipeline.

| input |  |  |  |  |  |  | continuity |  |  |  | comparison to truth data |  |  |  |  |  | run time |
| --- | --- | --- | --- | --- | --- | --- | --- | --- | --- | --- | --- | --- | --- | --- | --- | --- | --- |
| id | sample | ethnicity | sex | data description | X | F | N50 contig (kb) | N50 phase block (Mb) | N50 scaffold (Mb) | gappiness | N50 perfect stretch (kb) | phasing error rate | missing kmers (%) |  | inconsistent at given distance (%) |  | wall clock (days) |
|  |  |  |  |  |  |  |  |  |  |  |  |  | haploid | diploid | 1 Mb | 10 Mb |  |
| A | NA19238 | Yoruban | F | one 10x library | 56 | 115 | 114.6 | 8.0 | 18.7 | 2.1 |  |  | 14.5 | 10.0 | 1.2 | 0.8 | 1.7 |
| B | NA19240 | Yoruban | F | one 10x library | 56 | 125 | 118.8 | 9.3 | 16.4 | 2.3 |  | 0.00008 | 14.4 | 9.8 | 1.2 | 0.7 | 1.7 |
| C | HG00733 | Puerto Rican | F | one 10x library | 56 | 106 | 123.6 | 3.4 | 17.8 | 2.0 |  | 0.00008 | 12.7 | 9.2 | 1.0 | 1.2 | 1.7 |
| D | HG00512 | Chinese | M | one 10x library | 56 | 102 | 113.2 | 2.7 | 15.4 | 2.2 |  |  | 13.6 | 10.0 | 1.3 | 0.5 | 1.7 |
| E | NA24385 | Ashkenazi | M | one 10x library | 56 | 120 | 106.4 | 4.2 | 15.1 | 2.6 |  | 0.00006 | 13.9 | 9.6 | 1.3 | 2.0 | 1.8 |
| F | HGP | European | M | one 10x library | 56 | 139 | 120.2 | 4.5 | 18.6 | 2.5 | 19.8 |  | 12.4 | 8.8 | 1.8 | 0.9 | 2.0 |
| G | NA12878 | European | F | one 10x library | 56 | 92 | 118.5 | 2.8 | 16.4 | 2.0 | 16.5 | 0.00077 | 12.6 | 9.1 | 1.1 | 0.6 | 1.8 |
| H | NA12878 | European | F | unknown number of PacBio libraries plus BioNano Genomics data | 46 |  | 1594.2 |  | 25.4 | 4.6 |  |  | 18.0 |  | 0.5 | 2.0 |  |
| I | NA12878 | European | F | 6 libraries (fragment, jumping, 10x) | 160 |  | 12.3 |  | 30.1 | 10.2 |  |  | 19.7 |  | 1.1 | 7.1 |  |
| J | NA12878 | European | F | 9 libraries (fragment, jumping, Fosmid, Chicago) | 150 |  | 43.6 |  | 42.8 | 0.6 |  |  | 14.8 |  | 5.6 | 6.0 |  |
| K | NA24385 | Ashkenazi | M | 7 PacBio libraries | 71 |  | 4525.2 |  | 4.5 | 0.0 |  |  | 11.8 |  | 2.2 | 17.9 |  |
| L | NA24143 | Ashkenazi | F | 2 PacBio libraries | 30 |  | 1048.4 |  | 1.0 | 0.0 |  |  | 14.3 |  | 15.2 |  |  |
| M | YH | Chinese | M | ~18,000 Fosmid pools and 6 fragment and jumping libraries, Illumina sequenced, plus Complete Genomics data | 702 |  | 52.5 | 0.5 | 23.2 | 1.5 |  |  |  | 10.4 | 1.2 | 1.6 |  |

### *de novo* assembly

Because barcoded 10x data provide shallow coverage of each molecule, it is not possible to separately assemble the reads from each barcoded partition, which would otherwise be a natural approach (Voskoboynik et al. 2013). Instead the assembly process creates progressively larger contigs. At the point where many contigs are at least a few kb long, most molecules which ‘pass through’ a given contig will have at least one read landing on it. This information about partitions touching the contig may be used to link to other contigs, and moreover, all reads from all partitions touching the contig may be assembled together. This is the Supernova analog of single-partition assembly.

Following this strategy, barcodes play a relatively minor role in the initial process. To start out, we use a *de bruijn* graph approach (Pevzner et al. 2001), adapting the method of DISCOVAR (Weisenfeld et al. 2014). Kmers (K=48) are prefiltered to remove those present in only one barcode, thus reducing the incidence of false kmers, i.e. those absent from the sample. The remaining kmers are formed into an initial directed graph, in which edges represent unbranched DNA sequences, and abutting edges overlap by K-1 bases. Operations are then carried out to recover missing kmers and remove residual false kmers (Weisenfeld et al. 2014). At this point the graph (called the base graph) is an approximation to what would be obtained by collapsing the true sample genome sequence along identical 48-base sequences (Butler et al. 2008). We then use the read pairs to effectively increase K to about 200, so that the new graph represents an approximation to what would be obtained by collapsing the true sample genome sequence along identical 200-base sequences, thus achieving considerably greater resolution (**Methods**).

The remainder of the assembly process consists of a series of operations that modify this graph, so as to improve it. To facilitate these operations, we decompose the graph into units called lines (**Fig. 1**, **Methods**). Lines are extended linear regions, punctuated only by “bubbles”. Bubbles are places in the graph where the sequence diverges along alternate paths that then reconnect. Common sources of bubbles are loci that are heterozygous or difficult to read (in particular at long homopolymers).

**Figure 1.**
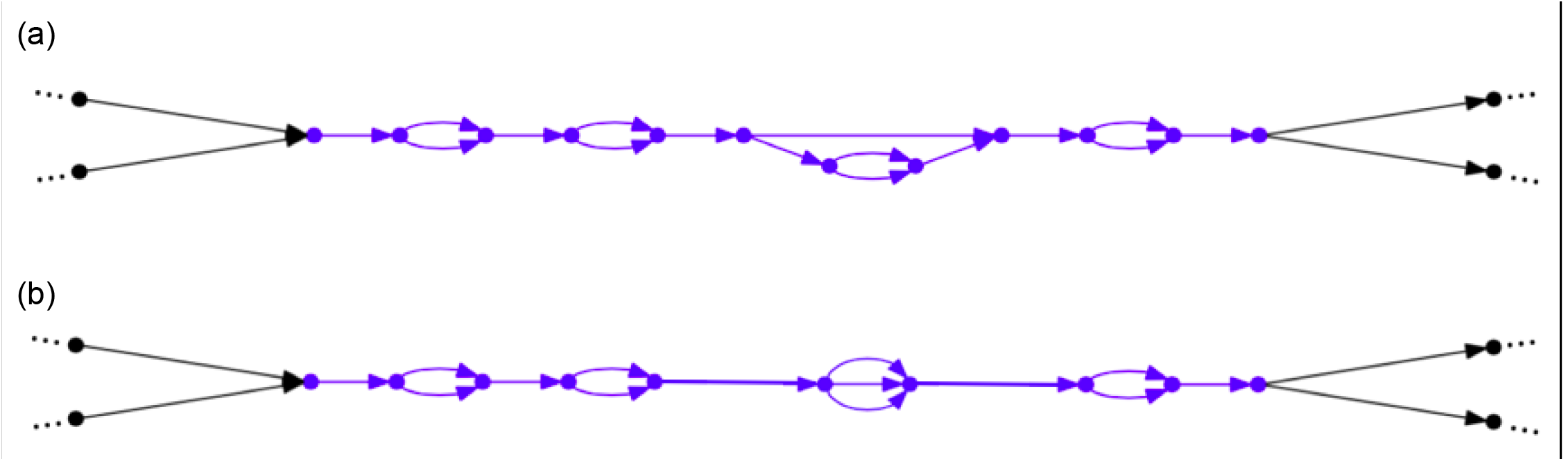
Lines in an assembly graph. Each edge represents a DNA sequence. (a) Blue portion describes a line in an assembly graph, which is an acyclic graph part bounded on both ends by single edges. The line alternates between five common segments and four bubbles, three of which have two branches. The third bubble is more complicated. The entire graph may be partitioned so that each of its edges lies in a unique line (allowing for degenerate cases, including single edge lines, and circles). (b) The same line, but now each bubble has been replaced by a bubble consisting of all its paths. After this change, each bubble consists only of parallel edges.

We can use lines to scaffold the assembly graph. This involves determining the relative order and orientation of two lines, then breaking the connections at their ends, then inserting a special ‘gap’ edge between the lines. The end result is a new line, which has a special ‘bubble’ consisting only of a gap edge. Subsequent operations (described later) may remove some of these gaps, replacing them by sequence.

Scaffolding is first carried out using read pairs. If the right end of one line is unambiguously connected by read pairs to the left end of another line, then they can be connected. Read pairs can reach over short gaps. To scaffold across larger gaps, we use the barcodes. Briefly, if two lines are actually near each other in the genome, then with high probability, multiple molecules (in the partitions) bridge the gap between the two lines. Therefore for any line, we may find candidate lines in its neighborhood by looking for other lines sharing many of the same barcodes. By scoring alternative orders and orientations (O&Os) of these lines, we can scaffold the lines by choosing their most probable configuration, excluding short lines whose position is uncertain (**Methods**).

Once the assembly has been scaffolded, some gaps may be replaced by one or more sequences. For short gaps, read pairs from both sides of the gap reach in and may cover the intervening sequence, from which it may be inferred. For long gaps, we first find the barcodes that are incident upon sequence proximate to the left and right sides of the gap. Then we find all the reads in these barcodes. This set of reads will include reads that properly lie within the gap, and yet be roughly ten times larger than that set (as each partition contains about ten molecules). We assemble this set of reads. Reads outside the gap locus tend to be at low coverage and hence not assemble. In this way it is typically possible to fill in the gap with a chunk of graph, and thereby remove the gap from the assembly. The chunk may not be a single sequence. For example at this stage heterozygous sites within the gap would typically be manifested as simple bubbles.

The final step in the assembly process is to phase lines. First for each line (**Fig. 1**) we find all its simple bubbles, i.e. bubbles having just two branches. Then we define a set of molecules. These are defined by series of reads from the same barcode, incident upon the line, and not having very large gaps (> 100 kb) between successive reads. A given molecule then ‘votes’ at certain bubbles, and the totality of this voting (across all molecules on each line) is then used to identify phaseable sections of the line, which are then separated into ‘megabubble’ arms (**Fig. 2**, **Methods**).

**Figure 2.**
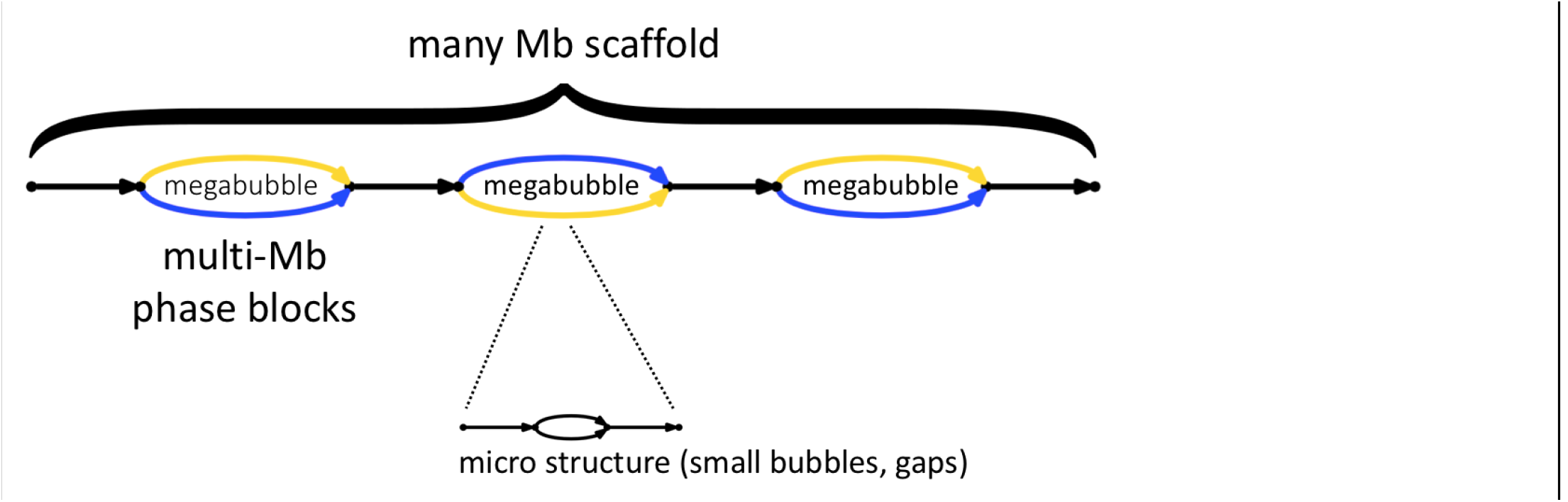
Supernova assemblies encode diploid genome architecture. Each edge represents a sequence. Blue represents one parental allele, gold the other. Megabubble arms represent alternative parental alleles at a given locus, whereas sequences between megabubbles are homozygous (or appear so to Supernova). Successive megabubbles are not phased relative to each other. Smaller scale features appears as gaps and bubbles.

### Software and computational performance

Supernova takes as input FASTQ files. No algorithmic parameters are supplied by the user. Supernova is designed to run on a single linux server. The peak memory usage across the seven human assemblies of this work was 335 GB, and accordingly we recommend use of a server having ≥ 384 GB RAM. Wall clock run times are shown in **Table 1**, and are in the range of two days.

### Supernova output

A Supernova assembly can separate homologous chromosomes over long distances, in this sense capturing the true biology of a diploid genome (**Fig. 2**). These separated alleles (or phase blocks) are represented as ‘megabubbles’ in the assembly, with each branch representing one parental allele. Sequences between megabubbles are nominally homozygous. Successive megabubbles are not phased relative to each other (if they were, they would have been combined). A chain of megabubbles as shown comprise a scaffold. In addition to large scale features, the Supernova graph encodes smaller features such as gaps and bubbles at long homopolymers whose length is not fully determined by the data.

A Supernova assembly can be translated into FASTA in several distinct ways, that might prove useful for different applications (**Fig. 3**). These allow representation of the full (or ‘raw’) graph (**Fig. 3a**), or erase micro features (choosing the most likely branch at small bubbles and replacing gap edges by Ns). There is more than one way to package the result, depending on how megabubble branch points are handled (**Fig. 3b-d**). We note that erasing micro features entails some loss of information, as in some cases the wrong branch of a bubble is chosen.

**Figure 3.**
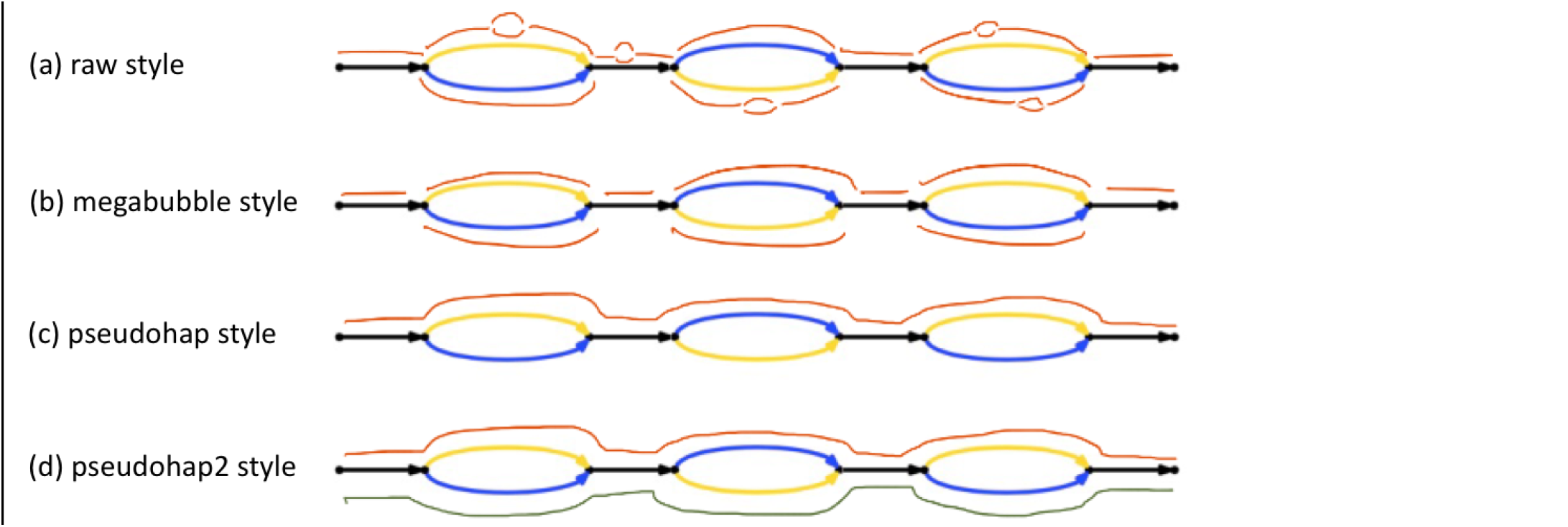
Representation of Supernova assemblies as FASTA. Several styles are depicted. **(a)** The raw style represents every edge in the assembly as a FASTA record (seen as red segments). These include microbubble arms and also gaps (printed as records comprising 100 Ns for gaps bridged by read pairs, or a larger number, the estimated gap size, **Supp. Note 5**). Unresolved cycles are replaced by a path through the cycle, followed by 10 Ns. Bubbles and gaps generally appear once per 10-20 kb, and consequently FASTA record from (a) are much shorter (~100 times) than those from (bcd). For each edge in the raw graph, there is also an edge written to the FASTA file representing the reverse complement sequence. For the remaining output styles, we flatten each microbubble by selecting the branch having highest coverage, merge gaps with adjacent sequences (leaving Ns), and drop reverse complement edges. **(b)** In this style each megabubble arm corresponds to a FASTA record, as does each intervening sequence. **(c)** The pseudohap style generates a single record per scaffold. As compared to the megabubble style, in the example one sees seven red edges on top (corresponding to seven FASTA records) that are combined into a single FASTA record in the pseudohap style. Megabubble arms are chosen arbitrarily so many records will mix maternal and paternal alleles. **(d)** This style is like the pseudohap option, except that for each scaffold, two ‘parallel’ pseudohaplotypes are created and placed in separate FASTA files.

Cycles in the graph provided an interesting test case. By a cycle we mean a set of one or more edges that include a path from a vertex back to itself. These are left intact in the full graph, however in the other forms are replaced by a path through the cycle that traverses each edge at least once, followed by Ns. This unfortunately signifies a gap (which could in principle represent any sequence), whereas the full graph indicates precisely which sequences could be present at the locus.

### Inferred DNA length

From each of the Supernova assemblies, we inferred the statistics of DNA molecules that were delivered to a partition and thence sequenced. This reflects the quality of input material as well as degradation during the initial steps of library construction. **Table 1** shows the inferred values for the length-weighted mean (LWM) of these molecules, as field **F**. It was in the range 92-139 kb. The HGP sample was obtained from fresh blood, and yielded the longest DNA. The other samples were obtained from cell lines. The sample NA12878 may have yielded the shortest DNA because of repeated handling of the DNA tube to create multiple libraries, as that DNA sample was used as a control for many experiments (not directly connected to this work).

### Assembly assessment

We assessed in the same way our seven human assemblies and six human assemblies from the literature that represent the state of the art, encompassing a wide range of laboratory approaches, from low coverage (30x) PacBio to complex combinations of multiple technologies at much higher coverage (**Table 1**). These comparison assemblies were downloaded from publicly accessible FTP sites (**Supp. Note 4**). For Supernova assemblies, we computed using the pseudohap FASTA output (**Fig. 3c**), except as noted.

To facilitate a uniform comparison, we computed all statistics from scratch (except as noted), rather than referring to published values. Before computing these statistics, we removed all scaffolds shorter than 10 kb from each assembly, thereby normalizing for differences in the actual cutoffs used in defining the assemblies, which would otherwise significantly affect statistics, including coverage of the genome.

To assess the continuity of the assemblies, we first computed the N50 contig size. The mean across the seven Supernova assemblies was 116 kb, with little variation. The three PacBio-based assemblies had much larger contigs, whereas contigs from the other assemblies were two-fold or more shorter than those from Supernova.

All of the Supernova assemblies were diploid, with N50 phase block size ranging from 2.7 to 9.3 Mb, with variability due presumably to varied ancestry and varied DNA length. Of the six other human assemblies, only the 702x assembly of YH (assembly M) was diploid, and it had an N50 phase block size of 0.5 Mb (as reported). The ~100 kb molecules underlying Linked-Reads enable the long phase blocks that are difficult to achieve with other technologies.

Scaffolds in the Supernova assemblies ranged from 15.1 to 18.7 Mb (N50). For the PacBio-only assemblies (KL), scaffolds are contigs, as these assemblies have no gaps; these scaffolds are much shorter than the Supernova scaffolds. The four combination assemblies (HIJM) had longer scaffolds, ranging from 23 to 43 Mb. The gappiness (fraction of Ns) in these scaffolds also varied greatly, from 0% for the PacBio-only assemblies, to ~2% for the Supernova assemblies, to ~10% for assembly I.

Any assessment of assembly continuity would be tempered by an assessment of the accuracy and completeness of those same assemblies. Although one could do this by comparing to a human reference sequence (and we do so later), the ideal would be to exploit ground truth data from the *same* sample that was assembled. These data would consist of clones (from individual haplotypes), that had been independently sequenced and assembled, and which were representative of the genome. We could find only two samples, for which such truth data was available and for which high quality DNA could be procured to create assemblies. These were the sample from a living Human Genome Project donor, which we refer to as HGP, for which 340 Mb of finished clones had been sequenced and assembled during the project, at great expense, and NA12878, for which we had previously sequenced and assembled 4 Mb of random clones (Weisenfeld et al. 2014). Although the HGP clones were not truly random, we reasoned that they comprised so much of the genome (~10%) that they would be reasonably representative of it. They comprise a remarkable and unique resource.

For a given sample, if we knew the exact sequence for each of its chromosomes, we could assess the accuracy of an assembly of that sample by enumerating maximal regions of the genome that are perfectly represented in the assembly. Such regions would be terminated by errors or gaps in the assembly. (Note that displaying the wrong allele would count as an error.) We call the N50 size of such perfectly represented regions the ‘N50 perfect stretch’. For diploid genomes, if one has both a diploid assembly (thus attempting to display all chromosomes), and representative finished sequence from the exact same sample (thus providing a sample of those chromosomes), then one can approximate the N50 perfect stretch (**Supp. Note 6**). There are no publicly available assemblies that satisfy these requirements, other than those generated by Supernova. In particular, we can assess the N50 perfect stretch for Supernova assemblies of HGP and NA12878 in **Table 1** (assemblies F and G). These were computed from the raw output (**Fig. 3**).

We found that the N50 perfect stretch in these Supernova assemblies was 19.8 kb for the HGP assembly, and 16.5 kb for the NA12878 assembly (**Table 1**). The difference might be attributable to sampling error as the finished sequence for NA12878 comprised 4 Mb whereas the finished sequence for HGP comprised 340 Mb. We further examined the the alignments of the finished sequence to the HGP assembly to understand the exact nature of the assembly defects that terminated perfect stretches. For example **Fig. 4** (and the corresponding alignments for thousands of other clones) show a preponderance of errors in low complexity sequence and in particular near long homopolymers. These errors might be attributable to library construction defects, sequencing defects, algorithmic defects, or possibly errors in the finished sequence (Lander et al. 2001).

**Figure 4.**
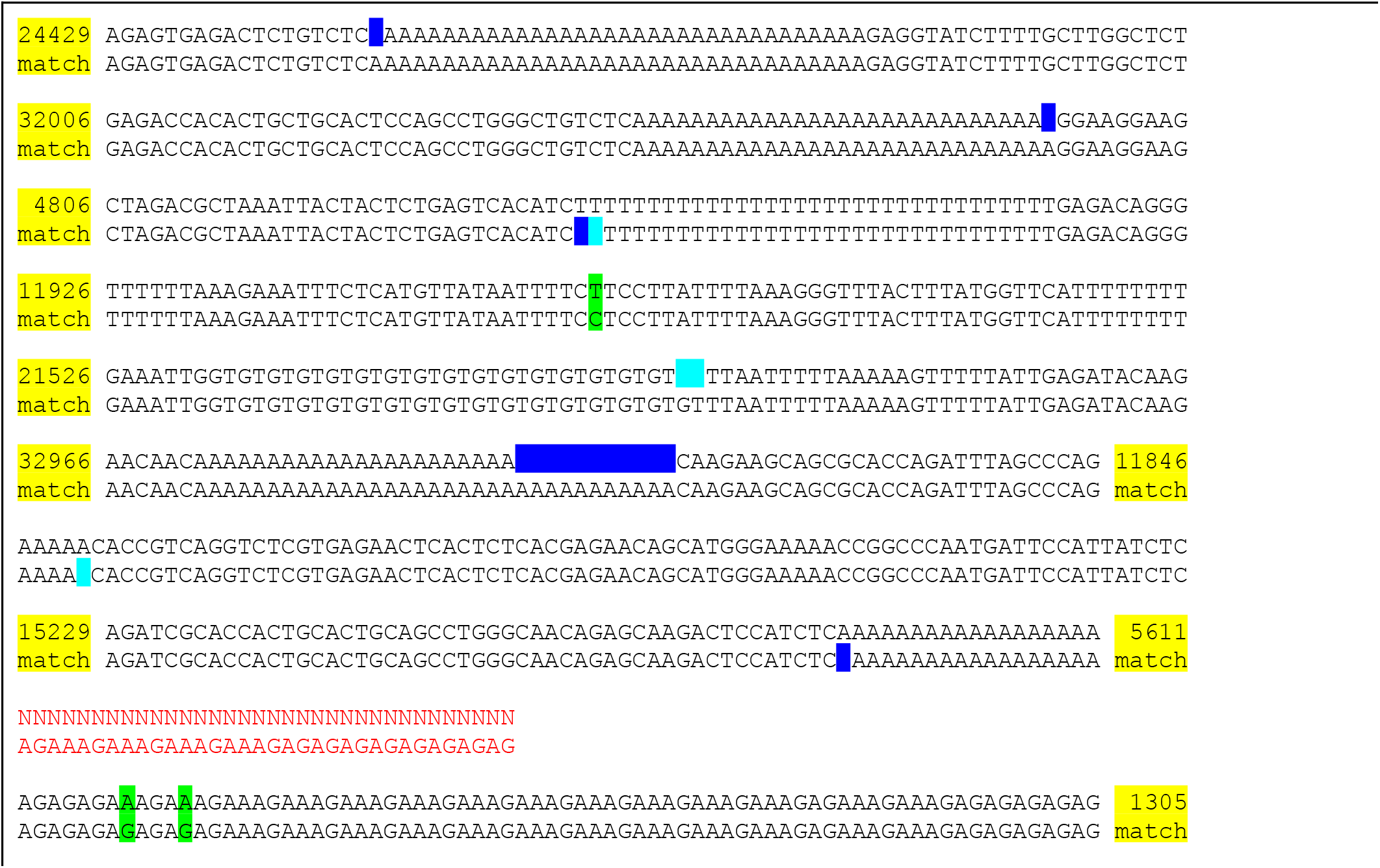
Alignment of Supernova assembly to finished sequence from the same sample. GenBank sequence AC004551.1 for finished clone RPCI1-71H24 has length 162,346 bases and its reverse complement perfectly matches GRCh37. The clone encompasses a region of Neandertal origin (Mendez et al. 2013). Both the clone and assembly F (**Table 1**) represent DNA from the same HGP donor. The clone matches a region of which 96% is between two megabubbles in the assembly, thus represented as homozygous. The alignment of the assembly to the clone region on GRCh37 is shown. Each line pair shows the assembly on top, reference on bottom. Yellow: abbreviated, perfectly matching stretches. Green: mismatched bases. Blue: indels. Cyan: indels, but not present in comparison to raw graph. Red: captured gap: signified by 34 Ns (actual number in assembly is 100); assembly region also has two cycles, each suffixed by 10 Ns in output, not shown. In these cases the flattened sequence for the cycle exactly matches the reference.

In more detail, **Fig. 4** displays the alignment of the Supernova HGP assembly to a 162 kb finished clone from the same sample, interesting because it is subsumes a region of Neandertal origin (Mendez et al. 2013). Each of several discrepancies between the assembly and finished sequence are annotated. Although we expect that most of these correspond to defects in the Supernova assembly, there is at least one case where the clone sequence is likely in error: for the first mismatch (shown in green), all reads in the Supernova assembly support the assembly base, and moreover the base was changed in the transition from GRCh37 to GRCh38, so that the latter now matches the assembly.

The exact list of discrepancies depends on the particular form of the Supernova assembly that the clone is compared to (**Fig. 4**), and this is interesting because it speaks to the power of the different forms. For this clone, only indel discrepancies are affected. If we compare the clone to the raw graph (full assembly), choosing the best path through the graph, then only the indel discrepancies shown in blue occur. However if we instead compare to the pseudohap style, in which small bubbles have been flattened by choosing the most probable path through the bubble (without looking at any reference sequence), then there are additional discrepancies, seen in cyan. Bubbles in the full assembly thus contain some information that is lost in the other forms.

Parental sequence data may also be used to assess the assembly of their child. In particular, this could provide a direct readout on the accuracy of phase blocks in a diploid assembly. This has not been done before for human genomes, because for the two extant diploid human assemblies (Levy et al. 2007; Cao et al. 2015), the parents were not sequenced. For four of the Supernova assemblies, (BCEG in **Table 1**), the parents had been sequenced, and phased VCFs were available (**Supp. Note 7**). This allowed us to estimate the phasing accuracy of these assemblies.

To do this, for each megabubble, whenever we found two positions on alternate megabubble branches that could be mapped to the same position on GRCh37, which represented different bases (a heterozygous SNP), and which was phased in the VCF, we recorded either 0 or 1, depending on whether the ‘top’ branch of the megabubble was assigned to the maternal or paternal allele. A sequence of all 0s or all 1s would represent perfect phasing. We counted all the ‘votes’ (0 or 1), and counted all ‘wrong votes’ (1 if majority = 0, 0 if majority=1), and summed across all megabubbles of size ≥ 1 Mb. The global error rate for phasing of a given assembly would be (wrong votes)/votes, noting that a ‘long switch’, i.e. haplotype misassembly error on even a single megabubble would drive this rate up.

We found that for three individuals (assemblies BCE), the observed phasing error rates were between 0.00006 and 0.00008, whereas for the the fourth individual (assembly G), the observed error rate was 0.00077. The difference is attributable to rare ‘long switch’ (haplotype misassembly) events, wherein maternal and paternal sequences are juxtaposed within a megabubble, and which just happened to have occurred in assembly G and not the others; for example exclusion of one such event lowers the rate to 0.00015.

Next, to measure the relative completeness of the different assemblies, we computed the fraction of nonduplicate kmers that were present in the reference sequence (GRCh37) and missing from a given assembly (**Supp. Note 8**). We used K = 100, balancing between two considerations. First it was important that the fraction of duplicate kmers be small, as the analysis would be blind to them. For K = 100, the fraction of duplicate kmers in GRCh37 is 2.3%. Second, we did not want to lose too many kmers to polymorphism. Assuming a polymorphism rate of 1/1000, we would expect about 10% of kmers would be missing because of *bona fide* differences between the sample and reference.

Most of the comparison assemblies were haploid and thus their missing fraction had to be computed based on their haploid output. For the Supernova assemblies, we computed both the haploid missing fraction (based on output type pseudohap) or their diploid missing fraction (based on output type pseudohap2). For assembly M (the 702x diploid assembly of YH), because there was no direct way to divide the assembly into haplotypes, we used the entire assembly and reported only the diploid missing fraction.

For the seven Supernova assemblies, the haploid missing fraction varied from 12.4 to 14.5%, with the highest values for the African samples (as would be expected, assuming that the African samples are most divergent from the reference). In general, the haploid coverage of the comparison assemblies is lower than that for the Supernova assemblies. For example, assembly I is missing 19.7%. The one exception is the 71x PacBio assembly of NA24385 (assembly K), which is missing 11.8%, as compared to the Supernova assembly of the same sample (assembly E), which is missing 13.9%. However, the corresponding *diploid* Supernova assembly is missing only 9.6%, again lower than the PacBio assembly (which is haploid).

We then assessed long-range accuracy of the assemblies. To do this, for a given assembly, and for fixed sizes (1 Mb, 10 Mb), we selected all scaffold segments (sequences of the given number of bases, within one FASTA record) of the given size in the assembly, whose end kmers occurred exactly once in the reference sequence. We excluded segments that bridged a gap of size 100 or more in the reference, as these gap sizes could be inaccurate or polymorphic (Bovee et al. 2008). For each segment, we tested its end kmers for consistent placement, meaning lying on the same chromosome, in the correct order and orientation, and defining a fragment whose length is within 10% of the fixed size. The fraction of segments whose end kmers were placed inconsistently is reported in **Table 1**. Inconsistency could be due to assembly error, or large errors in assembly gap measurement, or polymorphism within a sample or between it and the reference.

For the seven Supernova assemblies, and the two distances (1 Mb, 10 Mb), all inconsistent fractions were between 0.6% and 2.0%. Two of the comparison assemblies were comparably accurate: assembly H, based on PacBio and BioNano Genomics data, and assembly M (the 702x diploid assembly of YH). The other four comparison assemblies exhibited several-fold higher inconsistency at one or more measurement distances. For example, assembly J (including Dovetail data), had long-range inconsistencies of 5.6% and 6.0%, somewhat qualifying the advantage of its very long scaffolds (42.8 Mb). The 71x PacBio assembly (K) had an inconsistency of 17.9% at 10 Mb, suggesting that PacBio data alone *might* be insufficient for accurate long-range assembly.

## DISCUSSION

Although knowledge of the genome is a fundamental starting point for biology, for large and complex genomes, obtaining that knowledge continues to be a challenge. Low-cost and straightforward methods based on read alignment to a reference provide an extraordinarily valuable but incomplete readout. A far more complete picture can be obtained through complex and sophisticated *de novo* assembly approaches, but their material requirements and expense preclude widespread use.

Moreover, nearly all *de novo* assemblies of diploid genomes have been haploid: at each locus they combine together sequences from maternal and paternal chromosomes, yielding as output a single mélange. This both corrupts and loses information, and thus generating diploid assemblies has been a major goal of the field. It has been achieved for genomes up to 5% of the size of a human genome (Jones et al. 2004; Chin et al. 2016), and in two cases, at great expense, for human genomes (Levy et al. 2007; Cao et al. 2015).

In this work we demonstrate true diploid human assemblies, via a single straightforward library made from ~1 ng of high molecular weight DNA. We carried out our approach on seven human samples, which we sequenced on the Illumina HiSeq X instrument, at low cost. These assemblies used identical code, with identical parameters as a ‘pushbutton’ process that ran in two days on a single server. The aggregate experiment burden of our approach is dramatically lower than that for all of the human assemblies that we compared to. Our approach yields much longer phase blocks than the previous diploid human assemblies (Levy et al. 2007; Cao et al. 2015). Our diploid human assemblies are the first to be validated using finished sequence from the same sample, and the first whose phasing accuracy has been validated using parental sequences.

We anticipate utility of our new method both for routine use as a single-technology approach, and in combination with other technologies, e.g. for the ‘The Reference Genomes Improvement’ project (Steinberg et al. submitted). We have demonstrated our method here only on human genomes. Fundamentally different genomes (including much smaller ones, as well as polyploid genomes) will likely require modifications to our methods. However we expect that our methods will enable assembly of many similar genomes (including that of most vertebrates).

Our diploid assemblies open the door to new analytical approaches, including alignment of assemblies to a reference sequence to call variants. The low cost and burden of our approach makes it applicable to large scale projects, both for human and ‘new’ genomes, posing new opportunities and challenges both for experimental design and biological interpretation.

## METHODS

### Genomic DNA samples

Genomic DNA samples.Informed consent was obtained and peripheral blood samples were obtained from an anonymous donor (labeled HGP in **Table 1**). Genomic DNA (gDNA) was extracted from fresh whole blood according to 10x Sample Preparation Demonstrated Protocol “DNA Extraction from Whole Blood” (available at http://support.10xgenomics.com/de-novo-assembly/sample-prep/doc/demonstrated-protocol-hmw-dna-extraction-from-whole-blood). Optimal performance has been characterized on high molecular weight (HMW) gDNA with a mean length greater than 50 kb. The gDNA did not require further size selection or processing.

Commercially acquired samples were obtained (NA12878, NA19238, NA19240, HG00733, HG00512, NA24385, NA24143 from Coriell). Genomic DNA was extracted from human lymphocyte cells following the HMW gDNA extraction protocol outlined in the Chromium Genome User Guide Rev A (available at http://support.10xgenomics.com/de-novo-assembly/sampleprep/doc/user-guide-chromiumtm-genome-reagent-kit).

Genomic DNA was quantified with the Qubit dsDNA HS Assay Kit (Life Technologies) according to 10x Sample Preparation Demonstrated Protocol “High Molecular Weight DNA QC” (available at http://support.10xgenomics.com/de-novo-assembly/sample-prep/doc/demonstrated-protocol-hmw-dna-qc).

### Sequencing library construction using the Chromium Genome Reagent Kit

A Chromium Controller Instrument (10x Genomics, Pleasanton, CA) was used for sample preparation. The platform allows for the construction of 8 sequencing libraries by a single person in two days. Sample indexing and partition barcoded libraries were prepared using the Chromium Genome Reagent Kit (10x Genomics, Pleasanton, CA) according to manufacturer's protocols described in the Chromium Genome User Guide Rev A (available at http://support.10xgenomics.com/de-novo-assembly/sample-prep/doc/user-guide-chromiumtm-genome-reagent-kit). Briefly, in the microfluidic Genome Chip, a library of Genome Gel Beads was combined with an optimal amount of HMW template gDNA in Master Mix and partitioning oil to create GEMs. 1.25 ng of template gDNA was partitioned across approximately 1 million GEMs, with the exception of the peripheral blood sample which utilized 1 ng of template gDNA. Upon dissolution of the Genome Gel Bead in the GEM, primers containing (i) an lllumina R1 sequence (Read 1 sequencing primer), (ii) a 16 bp 10x Barcode, and (iii) a 6 bp random primer sequence were released. GEM reactions were isothermally incubated (30°C for 3 hours; 65°C for 10 min; held at 4°C), and barcoded fragments ranging from a few to several hundred base pairs were generated. After incubation, the GEMs were broken and the barcoded DNA was recovered. Silane and Solid Phase Reversible Immobilization (SPRI) beads were used to purify and size select the fragments for library preparation.

Standard library prep was performed according to manufacturer's instructions described in the Chromium Genome User Guide Rev A (available at: http://support.10xgenomics.com/de-novo-assembly/sample-prep/doc/user-guide-chromiumtm-genome-reagent-kit) to construct sample-indexed libraries using 10x Genomics adaptors. The final libraries contained the P5 and P7 primers used in lllumina bridge amplification. The barcode sequencing libraries were then quantified by qPCR (KAPA Biosystems Library Quantification Kit for Illumina platforms). Sequencing was conducted with an Illumina HiSeq X with 2x150 paired-end reads based on manufacturer's protocols.

### Internal representation of Supernova assemblies

An assembly is first represented as a directed graph in which each edge represents a single strand of DNA sequence. Abutting edges overlap by K-1 bases (K=48). This graph is called the base graph. Subsequently a new graph called the super graph is constructed in which each edge represents a path in the base graph. Each super graph edge may be translated into a DNA sequence. Where super graph edges abut, their associated sequences overlap by K-1 bases (K=48, the same as the base graph). An advantage of this approach was that it allowed for certain computations to be carried out on the base graph, once, so that experiments involving super graph algorithm improvement could be carried out, without repeating those computations. It is also a convenient mechanism for effectively changing the K value, without changing the actual K value in use.

### Generation of the initial super graph

For each read pair, where possible, we find one (or sometimes more) paths in the graph that could represent the sequence of the originating insert (Weisenfeld et al. 2014). These paths are represented as sequences of integers corresponding to the identifiers of edges in the base graph. Whenever there are two paths that perfectly overlap by K’=200 bases, we formally join them via an equivalence relation. This yields the super graph.

### Definition of lines

A line in a graph is an acyclic subgraph, having a unique source and sink (relative to the subgraph), with one edge exiting the source and one edge entering the sink (and maximal with respect to these properties). We also allow for the special case of circles, which are similar, but have no source or sink. Every edge lies in a unique line.

### Ordering and orienting lines using barcodes

For all lines in the assembly, we carry out an initial computation, which assigns a linear coordinate system to each line and marks the positions of uniquely placed reads on it, organized by barcode. Now for a given line set S, we score alternative O&O possibilities, as follows. Each O&O for S thus yields a sequence of barcoded read positions along a hypothetical merged line. We compute a penalty for the given O&O, which is a sum, over all its constituent barcodes. For each barcode, we first compute the mean separation between successive read placements for that barcode (in the merged line). Then we traverse these placements, in order, finding those pairs of consecutive placements that bridge a jump from one constituent line to another, and which may thus represent a misconnection. We divide the separation for this pair by the mean separation for the barcode. If the quotient is smaller than a fixed bound (2.0) we discard it on the theory that it is probably noise. The remaining quotient is added to the penalty (**Supp. Fig. 1**). A given O&O is treated as the ‘winner’ if its penalty is at least a fixed heuristic amount (60.0) less than that for competing tested O&O possibilities for the same set of lines.

### Phasing

A ‘phasing’ is an orientation of each bubble on a line, placing one of its branches on ‘top’ and the other on the ‘bottom’. Initially we choose an arbitrary orientation for the bubbles. Each molecule touches some bubbles, and thus (relative to a given phasing) may be represented as a sequence with entries +1 for top, −1 for bottom, or 0 for silent. A phasing is ‘good’ if each molecule is coherent, containing nearly all 1s or nearly all −1s (plus 0s at silent positions). Accordingly we define the score of a phasing to be the sum over all molecules of Max(pluses,minuses) - Min(pluses,minuses).

We then carry out iterative perturbations, each of which flips some bubbles, and keeping only those perturbations that increase the phasing score. Three types of perturbations are attempted:
1. We flip bubbles on a given molecule to make it completely coherent.
2. We flip an individual bubble.
3. We pivot at a given point, flipping all bubbles to its left.

This yields an initial phasing. We then look for weaknesses in it. First, if flipping a bubble has too small an effect on the score, we exclude it from the phasing operation. For example a bubble might arise at a long homopolymer whose length was fixed in the sample but changed during data generation. These uncertain bubbles are ‘copied’ to both megabubble branches. Second, if a pivot has too small an effect on the score, we break the phasing at the pivot point, yielding multiple phase blocks for the given scaffold. For example this could happen if a sufficiently long region in a given sample was homozygous.

## DATA ACCESS

We have created a temporary site at which the following materials can be accessed by reviewers of this work:

- The seven Supernova assemblies in **Table 1** (which are being deposited in GenBank).
- The reads that were used to create these assemblies (which are being deposited in the SRA).
- Instructions for downsampling to the exact same sets of 1200M reads (so as to exactly reproduce our results).
- A file of 3431 finished clones from GenBank for the HGP sample, that we used for assessment.
- Open-source code and executables for Supernova

### These will be made publicly available as soon as permanent sites are constructed

In addition, creation of a cell line from the HGP donor is in progress. The permanent location of all materials will be noted in the published version of this work.

## ACKNOWLEDGMENTS

We thank the entire team at 10x Genomics for developing the technology that enabled this project. We thank Rabbea Abbas, Brendan Galvin, Cassandra Jabara and Susanna Jett for preparing DNA and libraries. We thank Sofia Kyriazopoulou-Panagiotopoulou, Mike Schnall-Levin and Grace Zheng for comments on the manuscript. We thank Katie Kong for assistance in preparing the manuscript. The development of Supernova was greatly accelerated by access to genome assembly code developed at the Whitehead Institute Center for Genome Research and thence the Broad Institute during the period 2000-2015, and open sourced through github. This code was used to develop ARACHNE, ALLPATHS, ALLPATHSLG, DISCOVAR and DISCOVAR *de novo*. As the manuscript for the latter was unpublished, we note here our gratitude to NIH, Iain MacCallum, Ted Sharpe, and Eric Lander for their contributions to and support for that work.

